## supplementary Figures for "Mechanistic basis of the dynamic response of TWIK1 ionic selectivity to pH"

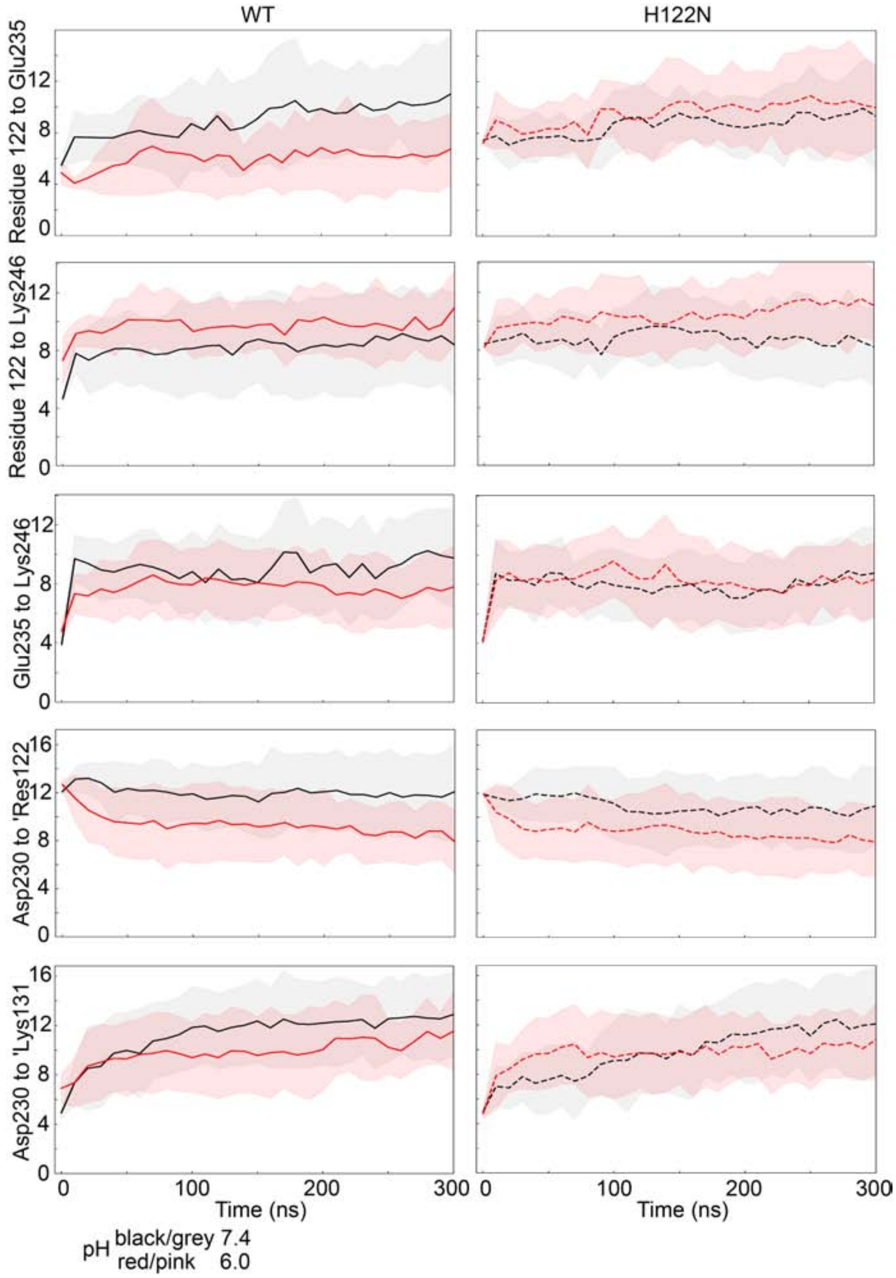


Figure S1. **Time series of distances between the centers of mass of side chain heavy atoms of pairs of residues.** Residues are shwon on the Y-axis, with distances involving neighboring subunits identified by the prefix '. The first 300 ns are shown, and distances were extracted every 10 ns. Lines and dotted lines represent averages, and shaded areas highlight standard deviations. Note that for dissociations of interacting residues (e.g. His122 and Glu235 at pH7.4), convergence is not expected.

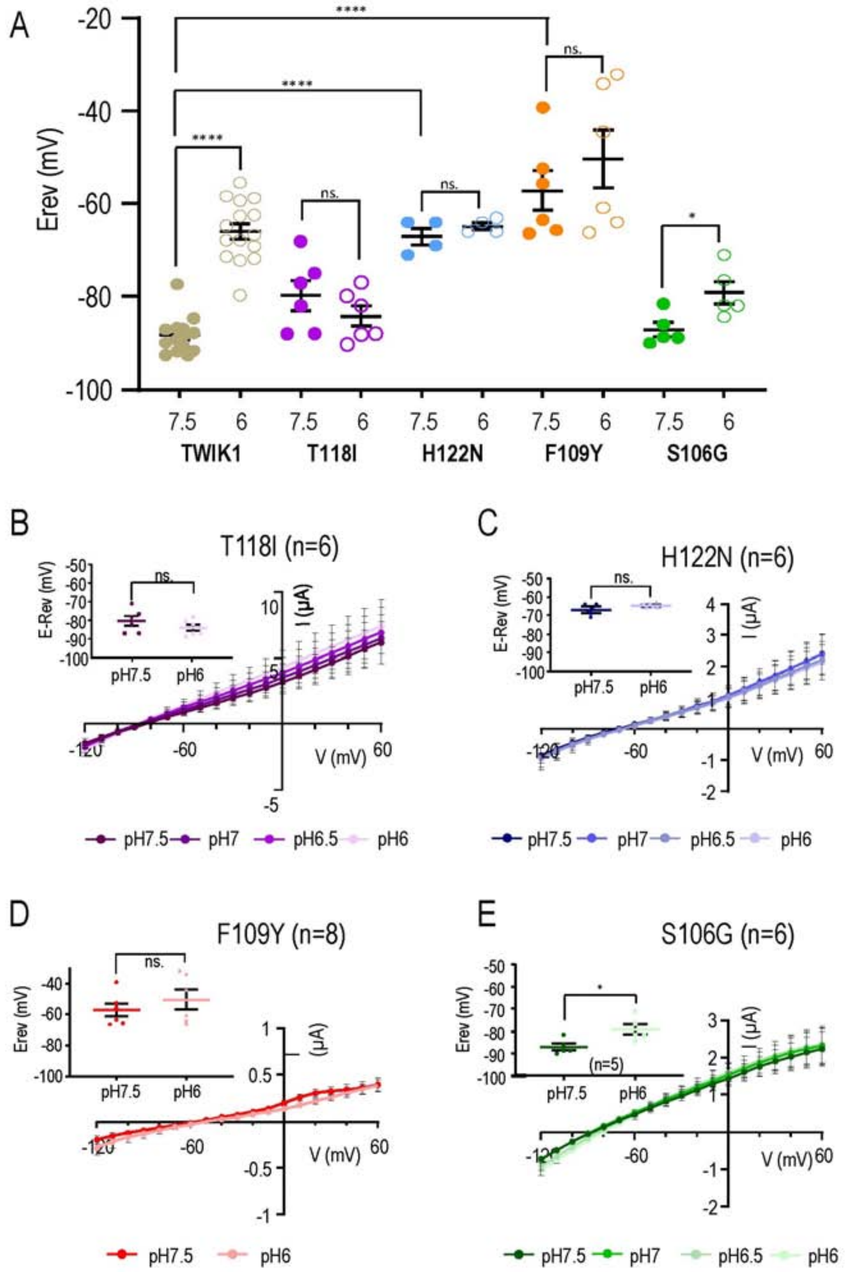


Figure S2. **pH-sensitivity of TWIK1 mutants.** (**A**) Reversal potential ($E_{Rev}$) variations between pH7.5 (solid dots) and pH6 (empty dots) of TWIK1 (brown), TWIK1.T188I (purple), TWIK1.H122N (blue), TWIK1.F109 (orange) and TWIK1.S106G (green). Values for TWIK1 are from Fig. 1B. Values for TWIK1 mutants T118I, H122N, F109Y and S106G are from Fig. S1. (**B-E**) Currents produced by TWIK1.T118I (**B**), TWIK1.H122N (**C**), TWIK1.F109Y (**D**) and TWIK1.S106G (**E**) in *Xenopus* oocytes at pH7.5, pH7, pH6.5 and pH6. After stabilization of the currents at pH7.5, voltage ramps were applied every ten seconds from -120 mV to +60 mV. pH was changed every 150 sec. Main panels: I/V curves obtained from stabilized currents at different pHs. Inset: reversal potentials ($E_{Rev}$) measured from stabilized currents at pH7.5 and pH6. All data are presented as mean ± SEM. The number of oocytes is indicated; ns, non-significant; *, $p < 0.05$; **, $p < 0.01$; ***, $p < 0.001$. ****, $p < 0.0001$.

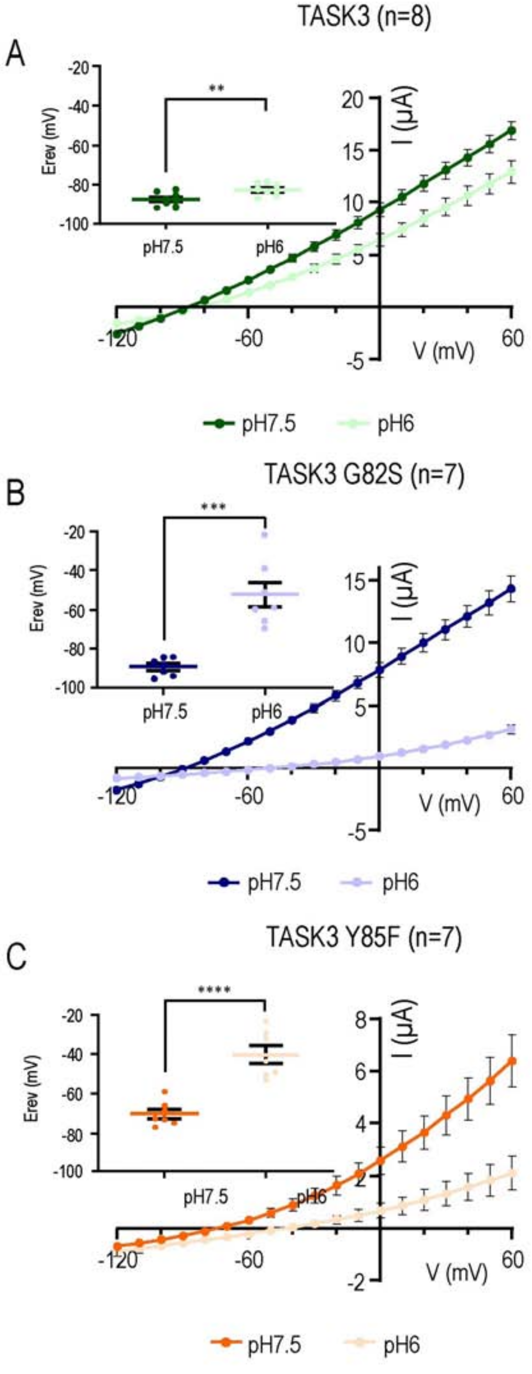


Figure S3. **pH response of currents produced by TASK3 and TASK3 mutants**. (**A-C**) Currents of TASK3 (**A**) and TASK3.G82S (**B**) and TASK3.Y85F (**C**) in *Xenopus* oocytes at pH7.5, Ph7, pH6.5 and pH6. Only currents recorded at pH7.5 and pH6 are shown. The number of oocytes is indicated. Potential ramps were applied from -120 mV to +60 mV. Inset: reversal potentials measured at pH7.5 and pH6; ns, non-significant; *, $p < 0.05$; **, $p < 0.01$; ***, $p < 0.001$. ****, $p < 0.0001$.

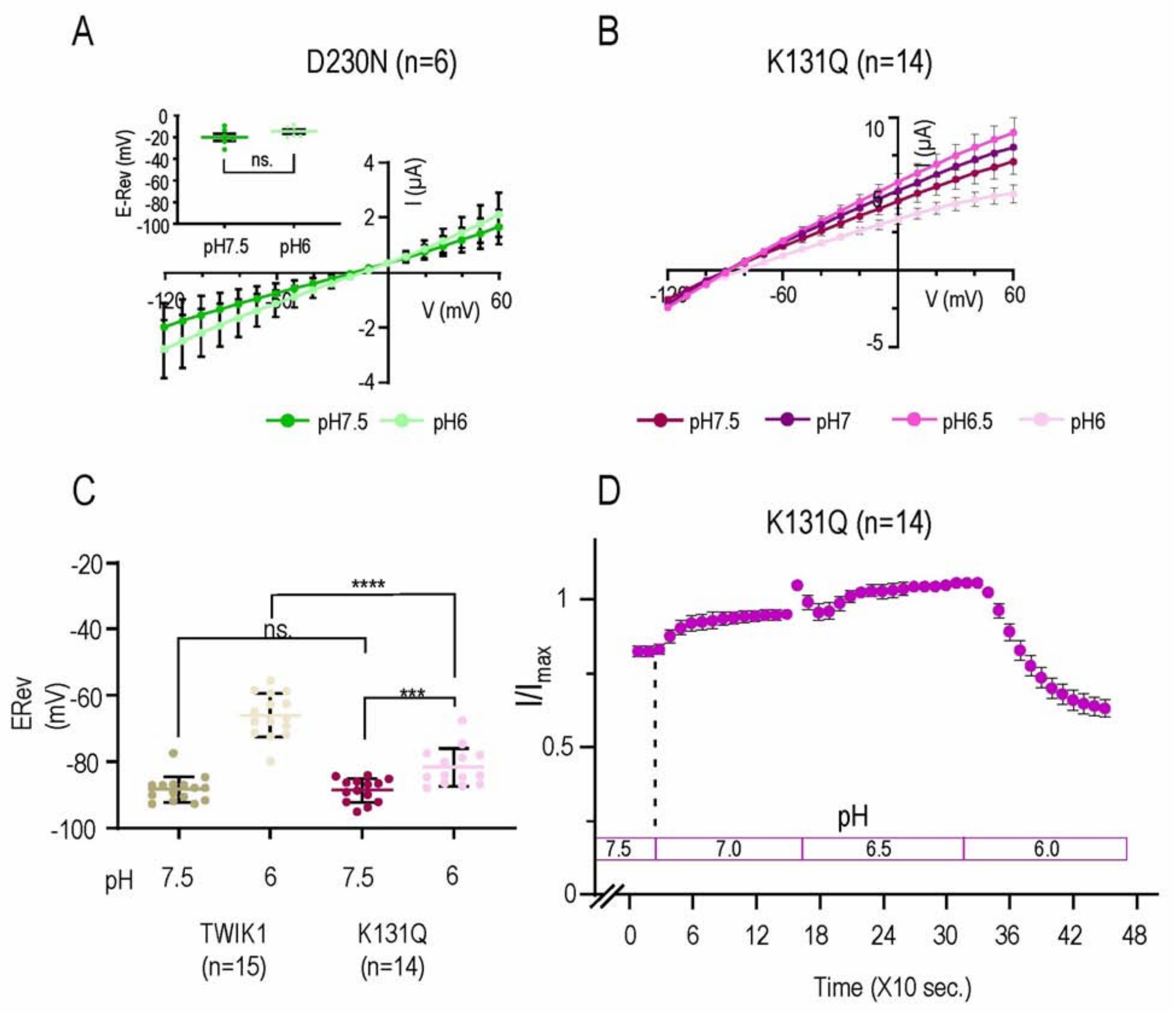


Figure S4. **pH-response of TWIK1.D230N and TWIK1.K131Q currents.** (**A, B**) Currents produced by TWIK1 mutants D230N (**A**) and K131Q (**B**) in *Xenopus* oocytes (the number of oocytes recorded is indicated). After current stabilization at pH7.5, voltage ramps were applied every ten seconds from -120 mV to +60 mV. For K131Q, pH was changed every 150 sec, starting from pH7.5 to pH6. Main panels: I/V curves recorded during the stabilized currents at different pH. Extracted from the voltage ramp recorded at each pH before changing to the next one. Inset: reversal potentials ($E_{Rev}$) of the stabilized currents at pH7.5 and pH6. (**C**) Reversal potentials ($E_{Rev}$) of TWIK1 (beige) and TWIK1.K131Q (purple) measured during the stabilized currents at pH7.5 and pH6 (in dark and clear respectively). Values for TWIK1 are from Fig. 1B. Values for TWIK1.K131Q are from Fig. S2B. (**D**) Kinetics of the TWIK1.K131Q current variations as a function of the pH, measured at 0 mV every 10 seconds. pH changes are indicated. All data are presented as mean ± SEM; ns, non-significant; *, $p < 0.05$; **, $p < 0.01$; ***, $p < 0.001$; ****, $p < 0.0001$.
